# A Computational Framework for Domain Insertion into Type IV Pili for Bacterial Display and Living Material Assembly

**DOI:** 10.64898/2026.06.09.731259.1

**Authors:** Robert F. Tesoriero, Nicole E. Harris, Olivia D. Suggs, Caroline M. Ajo-Franklin

**Affiliations:** Department of BioSciences, Rice University, Houston, TX, USA; Department of Bioengineering, Rice University, Houston, TX, USA; Department of Chemical and Biomolecular Engineering, Rice University, Houston, TX, USA; Rice Synthetic Biology Institute, Rice University, Houston, TX, USA

## Abstract

Bacterial surface structures have enabled display systems with broad impact across biotechnology, but their narrow host range limits their deployment into diverse species. Conversely, Type IV Pili (TFP) are ubiquitous, structurally conserved appendages found across bacteria, but have been minimally explored for display. Here, we describe a computational framework for predicting viable insertion sites in major pilins for stable TFP-mediated display, which we apply to the major pilin PilA1 of the cyanobacterium *Synechocystis* sp. PCC 6803 to enable covalent binding to living materials. By analyzing an Alphafold3-generated PilA1 monomer alongside known multimeric TFP multimers, our pipeline identifies non-interfacial, solvent-accessible, and flexible sites for optimal PilA1 display. We probe these sites with both full-length and truncated SpyCatcher003 at two different expression levels. We show that cells expressing these PilA1-SpyCatcher003 fusions maintain up to 8-fold higher levels of cell suspension than previous C-terminal PilA1 display platforms, suggesting improved TFP assembly despite more than a two-fold increase in cargo size. Additionally, we validate SpyCatcher003 reactivity across the engineered strains, enabling covalent attachment of SpyTag003-containing proteins on the *Synechocystis* surface. Lastly, we utilize this covalent patterning to achieve a four-fold increase in *Synechocystis* loading into a living material without compromising its viscoelastic or mechanical properties. Taken together, this work provides a predictive framework for TFP engineering, and opens the door towards programmable surface display across the breadth of bacterial species.

## Introduction

Bacterial surface structures, including curli, flagella, and surface layers mediate critical interactions between cells and their environments.^1–4^ These structures possess both high surface accessibility and density, which make them ideal scaffolds for displaying functional peptides and proteins.^1,2,4^ To date, surface structure-mediated bacterial display has achieved far-reaching impacts across fields^5–13^ from bioremediation^9^ and vaccines^8,14^ to biocatalysis^15^ and biomaterials.^5,11,16^ Extending these display systems into non-model species would enable deployment into diverse environments from hydrothermal vents^17^ to the human body,^18^ and the creation of patterned, carbon-neutral living materials. However, current display platforms are constrained to specific organisms, phylogenies and environments. To overcome these barriers, there is a need for generalizable strategies that expand structural display capacity across the bacterial kingdom.

Type IV Pili (TFP) are ubiquitous bacterial surface structures found in 30-45% of bacterial species across nearly all classes and environments.^19,20,18^ The roles of TFP are almost as wide ranging as their distribution, performing diverse functions such as motility, surface adhesion, biofilm formation, and DNA uptake.^20–22^ Key to this multifunctionality is the TFP’s structure, which is composed of monomers called pilins. The major pilins, which comprise the bulk of the fiber, possess a highly homologous structure despite large sequence diversity.^19,22^ These major pilins, together with functionally specialized minor pilins, are assembled into dynamic and retractable TFP fibers that adapt to the cell’s current needs.^21^ This combination of ubiquity and versatility make TFP a promising system for bacterial display, with potential for broad utility across chassis and applications.

While TFP have been engineered to display peptides and proteins, existing methods possess significant limitations. Cengic *et al*. engineered the C-terminus of the major pilin PilA1 from the cyanobacterium *Synechocystis* PCC 6803 (hereafter referred to as *Synechocystis*) to display the 6.5 kDa affibody Z_Taq_.^23^ However, Z_Taq_ display significantly impacted TFP assembly, leading to truncated TFP and limited display capacity. Inserting a smaller 1.8 kDa gold-binding peptide in the same PilA1 location permitted display and TFP assembly,^24^ demonstrating stable TFP display of a peptide. However, the ability to display entire proteins or protein domains in TFP without inhibiting assembly remains an open challenge.

The recent explosion of computational tools for protein structure prediction provides an excellent opportunity for improving TFP display.^25^ Machine learning models such as Alphafold2 can accurately predict protein structures from sequence alone, and are particularly effective in predicting highly conserved structures like TFP major pilin monomers.^26,27^ However, even Alphafold3, the current gold standard in protein structure prediction at time of writing, fails to accurately predict large assemblies like TFP multimers, frequently orienting pilin monomers within a fiber incorrectly or failing to produce fibrillar structures at all.^28,27^ This issue is fueled by a dearth of available multimeric TFP assemblies, with less than 10 solved structures available.^29–34^ Additionally, there have been limited examples utilizing these tools to predict structures that possess heterologous domain insertions, making computational design of TFP-mediated display difficult.

To overcome the current challenges of TFP-mediated display, we have developed a computational framework for predicting stable domain insertion sites into TFP, using Synechocystis PilA1 as a test case. We show that our predicted sites enable the functional display of a protein domain, SpyCatcher003, on the cell surface with minimal impacts to Type IV pilus assembly. Finally, we demonstrate the utility of TFP-mediated SpyCatcher003 display in living materials by enabling covalent SpyCatcher-SpyTag crosslinking of *Synechocystis* cells into a living structure.

## Results

### A computational pipeline for selecting insertion sites into the *Synechocystis* major pilin PilA1

Type IV pili assembly into micro-scale fibrillar structures relies upon precise interactions both within and between pilin monomers. Preserving these interactions is crucial for stable domain insertion.^21^ Additionally, any potential insertion site must be surface accessible and flexible to ensure proper display.^35^ Therefore, we reasoned that a successful insertion must: i) minimize disruptions to multimeric pilus assembly, ii) maximize surface accessibility of the insert, and iii) minimize local disruptions to monomer folding. To computationally predict insertion sites that meet these criteria, we performed three major structural analyses of PilA1: we 1) predicted intermolecular chain-chain interfaces to avoid disrupting of multimeric pilus assembly (criteria i), 2) calculated the solvent accessible surface area (SASA) of each residue to maximize insert accessibility (criteria ii), and 3) identified flexible regions within PilA1 to avoid impeding monomer folding (criteria iii) (Fig. 1A). We hypothesized that a computational pipeline combining these three analyses would lead us to domain insertion sites capable of stable TFP-mediated display.

**Figure 1.**
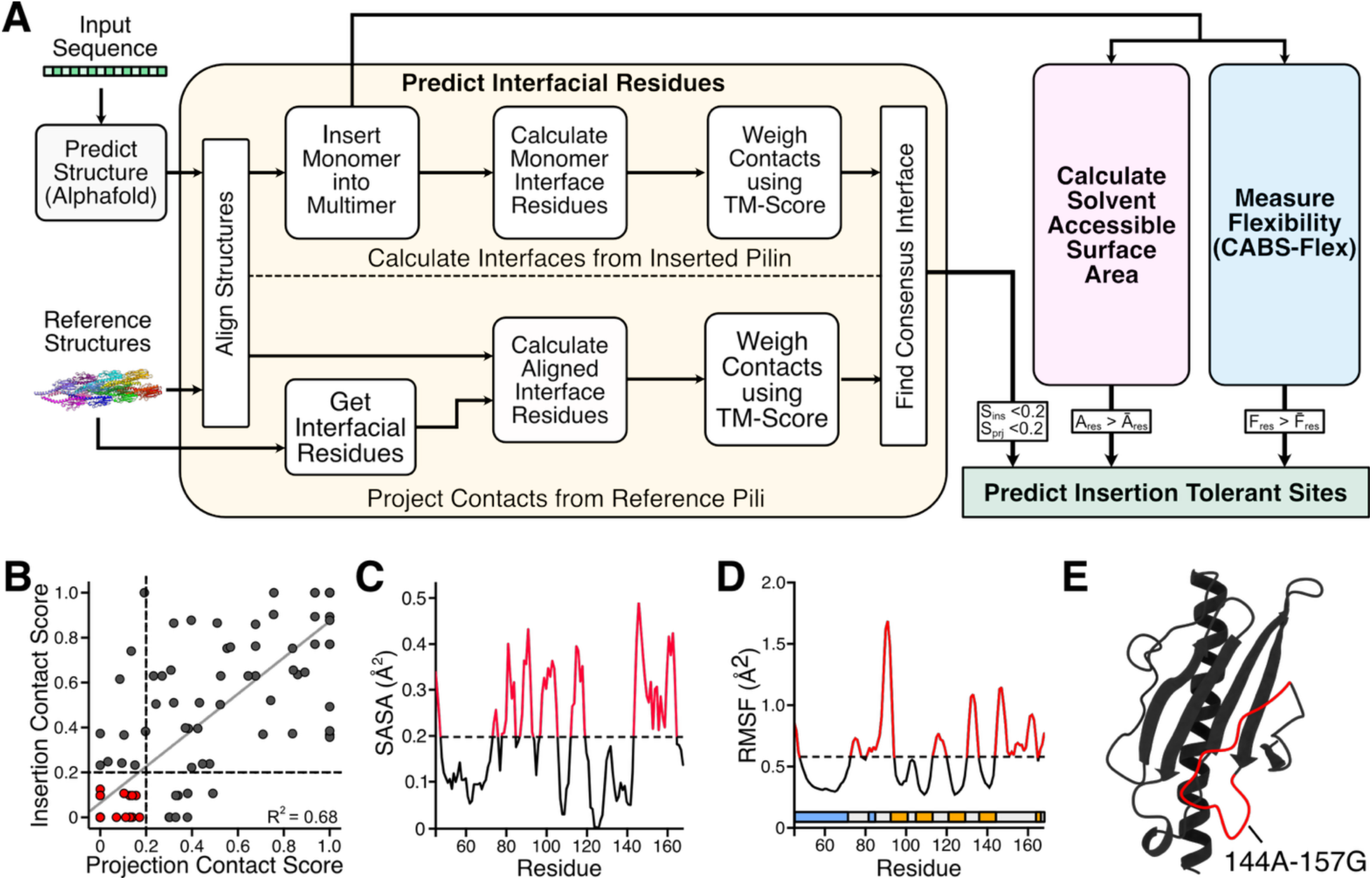
Computational pipeline for predicting insertion sites in the major pilin PilA1 of *Synechocystis* PCC 6803. **A)** Algorithm for predicting insertion tolerant sites of the pilin PilA1 using the sequence of PiA1 and reference multimeric Type IV pili structures. Interfaces are both mapped onto PilA1 from reference structures and calculated by inserting PilA1 into each multimer to produce a consensus interface. This information is combined with the solvent accessible surface area and flexible regions to determine insertion tolerant sites. **B)** Predicted consensus interface residues in PilA1, as determined by both inserting the PilA1 monomer into reference pilus structures and projecting native pilus interfaces onto PilA1. Only residues that were predicted as non-interfacial by both methods, labeled in red, were considered as potentially viable for insertion. **C)** Relative Solvent Accessible Surface Area (SASA) as a function of residue number, as determined by Shrake-Rupley calculations on PilA1 monomers inserted into reference pili. Each point represents a moving average of the two residues before and after each site. Red regions represent above average SASA measurements. **D)** Measurement of root mean square flexibility for each residue of PilA1, measured by CABS-FLEX. Each measurement represents a moving average of the two residues before and after each site. Red regions represent local maxima. Bar underneath plot maps to PilA1 secondary structure, with blue representing *ɑ*-helices, orange representing ꞵ-sheets, and white representing loops. **E)** Predicted insertion tolerant sites (red loop) in PilA1 (black), that represent flexible, solvent accessible regions not predicted to exist at chain-chain interfaces.

To build this pipeline, we first required structural information on both the monomeric and multimeric forms of PilA1. Given that the structure of PilA1 has not been solved, we first generated a high-confidence PilA1 monomer using Alphafold3 (Supplementary Figure 1).^27^ However, Alphafold3 failed to predict accurate PilA1 multimers (Supplementary Figure 2). To overcome this issue, we compiled a reference set of all multimeric TFP structures available in the RCSB Protein Data Bank (Supplementary Table 1). Comparing PilA1 to this reference set revealed that monomeric PilA1 possessed a single long N-terminal helical domain, whereas the multimeric reference structures possessed two shorter helices over the same region (Supplementary Figure 3). This difference is known to occur during the TFP multimerization process, where the N-terminal helix partially unfolds at a highly-conserved proline residue at position 22 (Supplementary Figure 4).^21^ We anticipated that these differences might lead to errors in structural alignment between PilA1 and known TFP, and so all structures were trimmed through residue Pro22 prior to our structural analysis pipeline.

With a monomeric and multimer structure of PilA1 in hand, the first stage of our pipeline was to predict which PilA1 residues participate in intermolecular chain-chain interfaces. To maximally leverage our small TFP dataset while minimizing errors, PilA1 interfacial residues were calculated by two methods. In the first method, we modeled the PilA1 multimer by systematically replacing a central chain in of the seven-member reference set with a trimmed PilA1 monomer via Template Modeling (TM)-Align. For each model, we used a K-nearest neighbors algorithm to calculate interface residues, defined as two atoms on separate polypeptides within 4.5 Å of each other. Residues possessing at least one contact were given an interface score of 1; all other residues were considered non-interfacial and given a score of 0. For each residue we then calculated a weighted average of these binary scores, weighing each by its respective TM-score between the reference TFP and PilA1 (normalized to PilA1 length). This calculation yielded a normalized set of insertion interface scores (Figure 1B).

Following this first calculation of interface residues, we reasoned that gaps in the alignment between PilA1 and the reference TFP structures may introduce interface prediction errors. We hypothesized that mapping the native interface residues from the reference TFP structures onto PilA1 would provide a complementary second interface measurement. Reference TFP interfaces were first calculated with an interchain atom distance of 4.5 Å and mapped to PilA1 using the TM-alignments. These mapped residues were then weighted via TM-score and normalized residue-wise to produce mapped interface scores. This second prediction showed a good positive linear correlation with the first (Figure 1B, R^2^ = 0.68). To obtain a final prediction, we combined the two score sets, considering any residue with a prediction score greater than 0.20 from both sets as interfacial (Figure 1B). As expected, most interfacial residues were predicted to occur in the N-terminal helix and the periphery of the pilin body, demonstrating the accuracy of our method.

Next, we determined solvent accessible residues by calculating the solvent accessible surface area (SASA) of the TFP-inserted PilA1 structures using the Shrake-Rupley algorithm. Each set of SASA measurements were weighted by their respective TM-scores (Figure 1C).^36^ To capture local accessibility across the structure, we utilized a sliding window approach, where the SASA values of the two residues before and after each insertion site were averaged. To filter out the most inaccessible regions while retaining as many sites as possible, all sites possessing SASA values above this average (0.200 Å^2^) were considered solvent accessible. The N-terminal domain was largely inaccessible, but there were several accessible regions across the middle of the structure near the first β-sheet and the C-terminal loop.

Our third structural analysis was to identify flexible regions within our multimeric PilA1 models. We calculated the root mean squared flexibility (RMSF) of the reference TFP-inserted PilA1 monomer. RMSF was measured using CABS-Flex in Category mode, which leverages the predicted local difference distance test (pLDDT) metric of the predicted structure to improve measurement accuracy (Figure 1D).^37^ Similar to the SASA measurements, RMSF was weighed by TM-score and averaged using a 6-residue sliding window, yielding an average RMSF value of 0.589 Å^2^. Residues with above average flexibility, labelled in red, generally corresponded to unstructured regions.

To generate a final prediction of insertion sites, the interface prediction, SASA, and RMSF were combined such that only sites that passed all three sets of binary thresholds were considered as insertion-tolerant. These sites were largely localized within the C-terminal loop region, with isolated sites in other loop regions (Supplementary Figure 5). The largest contiguous region, residues 144G-157G in the C-terminal loop, was chosen as the optimal target for insertion (Figure 1E). To cover this area effectively, we selected 3 predicted insertion-tolerant sites: 147G:148S, 150T:151P, and 156G:157G. Additionally, we wanted to demonstrate the ability of our pipeline to correctly determine sites that could not tolerate insertion. As such, we also selected 133G:134T, which was predicted to have high flexibility and high probability of being located at a chain-chain interface. These four sites, hereafter referred to as 133G, 147G, 150T, and 156G were chosen to probe whether insertions could be tolerated in PilA1 without disrupting assembly.

### Design of Expression Systems for PilA1-SpyCatcher003 Fusion proteins

Beyond site insertion tolerance, both the protein expression level and cargo size are critical parameters for successful TFP display.^38–40^ To understand how these parameters impact PilA1-mediated SpyCatcher display, we designed a set of 4 constructs for each of the 133G, 147G, 150T, and 156G insertion sites, varying both expression level and cargo size (Supplementary Figure 6, Supplementary Tables 3, 4). To vary expression level, we selected two constitutive synthetic promoters, s7 and s6, designed by Seo *et al*. to express at low and medium levels, respectively.^41^ For clarity, we will hereafter refer to these promoters by their expression level, namely ‘Low’ (L), and ‘Medium’ (M). Likewise, to probe the impact of cargo size on TFP display, we designed two SpyCatcher003 variants of different molecular weight. Prior work has shown that the 21 N-terminal and 10 C-terminal residues of SpyCatcher are disordered, and that removing these regions yields a ∼25% decrease in molecular weight while minimally impacting reactivity.^42^ Thus, we selected this truncated 82 residue mutant (9.08 kDa), dubbed SpyCatcher003ΔNΔC, as our small insert, and the full-length 110 residue (12.13 kDa) SpyCatcher003 as our large insert. Furthermore, to ensure minimal interactions between PilA1 and SpyCatcher003, all inserts were flanked on both sides by a 12-residue (GGSG)_3_ motif. The sixteen combinations of promoters, insertion sites, and insertion sizes were constructed into a RSF1010-based vector by Golden Gate assembly, sequence verified and transformed into *Synechocystis* via electroporation. These constructs were expressed alongside native PilA1 expression, as described previously.^23^ Strains are hereafter designated by expression level, insertion site, and SpyCatcher003 size (e.g., ‘L-147G:SC3ΔNΔC’ denotes SpyCatcher003ΔNΔC inserted into PilA1 site 147G under the low-expression s7 promoter).

### TFP assembly is influenced by SpyCatcher003 size and insertion site

In aqueous environments, TFP help bacterial cells remain in suspension, while cells lacking TFP sediment more quickly because of their altered surface area and charge.^43,44^ Therefore, defects in TFP assembly can be detected by monitoring changes in cell suspension density over time. To characterize TFP assembly in our PilA1-SpyCatcher003 displaying strains, we monitored the cell density in the uppermost ∼5 millimeters of cell suspension using OD_730_ over 16 hours (Supplementary Figure 7). After 16 hours, all strains displaying SpyCatcher003ΔNΔC retained approximately 80% of cells in suspension, regardless of insertion site (Supplementary Figure 8). In contrast, strains displaying full-length SpyCatcher003 exhibited strong insertion site-dependent sedimentation, with only sites 147G and 156G maintaining over 50% suspension (dotted line) at both expression levels (Figure 2A).

**Figure 2.**
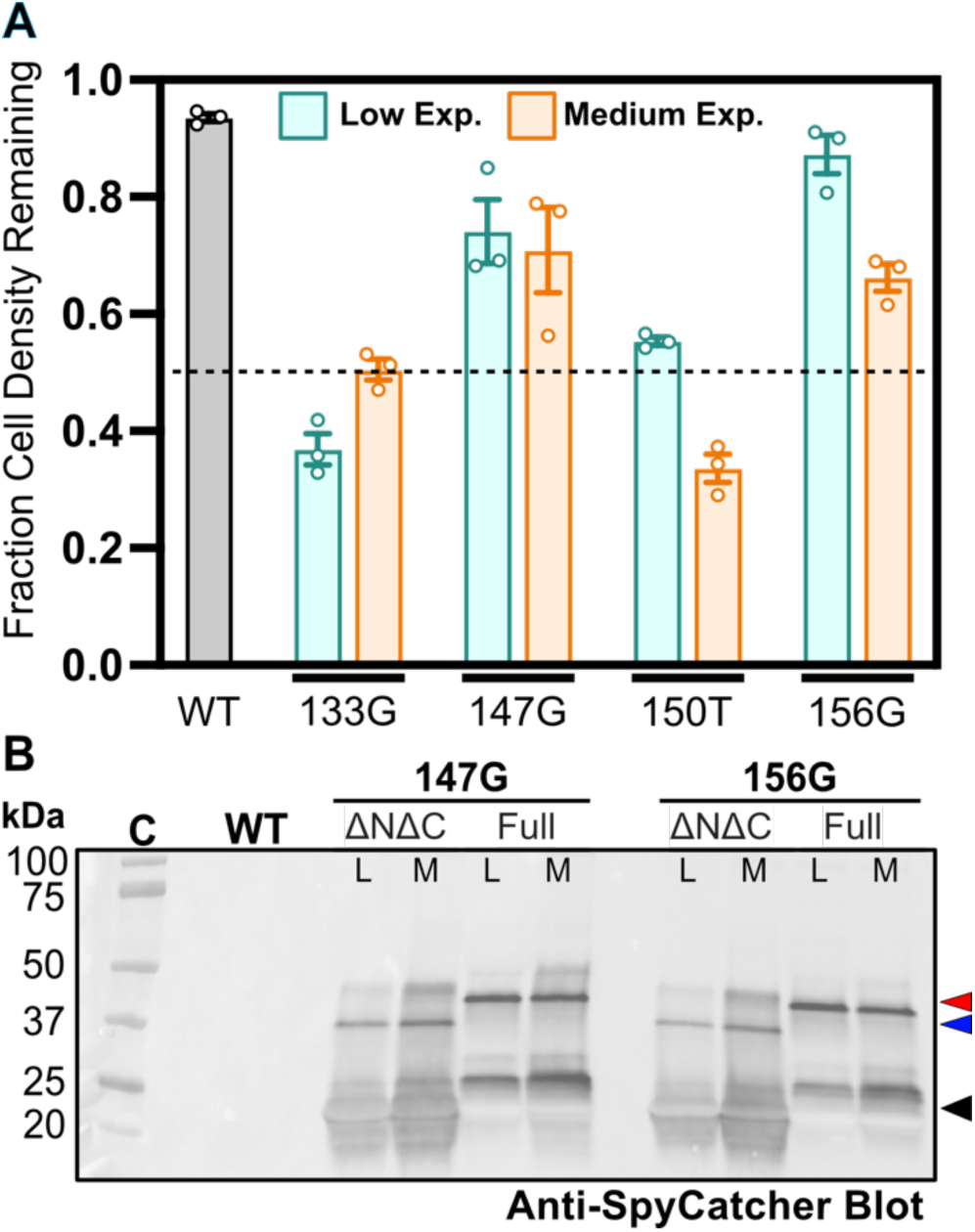
Expression and Display of PilA1-SpyCatcher003 impacts cell sedimentation in an insertion site-dependent manner. **A)** Remaining cell density in the top layer of different *Synechocystis* strains in solution after 16 hr, demonstrating insertion site and expression level dependent sedimentation. The black dotted line signifies 50% remaining cell density. **B)** Anti-SpyCatcher western blot of *Synechocystis* surface proteins, showing the display of full-length PilA1-SpyCatcher003. Red, blue and black arrows indicate PilA1-SpyCatcher003, PilA1-SpyCatcher003ΔNΔC, and wild-type PilA1, respectively.

A three-way ANOVA revealed that insertion site was the most influential parameter for sedimentation, both in isolation and when interacting with cargo size and expression level (Supplementary Table 5). The interaction between insertion site and cargo size was particularly distinct, highlighting how site selection is critical for maintaining display capacity. Overall, 147G and 156G, retained closest to wild-type suspension across all conditions, suggesting the smallest defect to pilus assembly. These strains also exhibited little to no growth defects (Supplementary Figure 9, Supplementary Table 6), and so were selected for further characterization.

To confirm that the 147G and 156G strains displayed PilA1-SpyCatcher003, TFP were sheared and extracted from the *Synechocystis* surface,^23^ then analyzed by SDS-PAGE (Supplementary Figure 10) and anti-SpyCatcher western blot (Figure 2B). SDS-PAGE analysis revealed a strong band for each sample at 22 kDa corresponding to the wild-type, glycosylated PilA1 monomer.^23,45,46^ Surprisingly, no distinct bands were observed at expected molecular weights for the unglycosylated or glycosylated fusion pilins (25.9 kDa and 39 kDa for PilA1-SpyCatcher003ΔNΔC; 28.9 kDa and 44 kDa for PilA1-SpyCatcher003, respectively), suggesting low abundance of of fusion pilins within the pilus fiber.

To more sensitively detect PilA1-SpyCatcher003 incorporation, we probed samples by anti-SpyCatcher Western blot. Multiple bands were observed across all engineered strains. Strains displaying SpyCatcher003ΔNΔC showed bands at 25.9 kDa and 39 kDa. Likewise, strains displaying full SpyCatcher003 exhibited bands at 28.9 kDa and 44 kDa. For both sets of strains, these bands corresponded to unglycosylated and glycosylated fusion PilA1 respectively, suggesting incomplete glycosylation of the fusion pilin pool. Nevertheless, the presence of intact PilA1-SpyCatcher003 on the cell surface of the 147G and 156G strains validated our prediction that these sites could tolerate insertions.

### PilA1-SpyCatcher003 displaying strains exhibit insertion site-dependent reactivity

Having confirmed the surface display of PilA1-SpyCatcher003, we evaluated if the inserted cargo was functional. We probed SpyCatcher003 reactivity by incubating the WT, 147G, and 156G strains with SpyTag003-sfGFP-HisTag for 6 hours, washing the cells, and measuring sfGFP fluorescence. SpyTag003-sfGFP binding was first quantified by flow cytometry (Supplementary Figures 11,12). Fluorescence intensity values from two independent trials were normalized to the median wild-type fluorescence.

With the exception of the L-147G:SC3 strain, which exhibited a 1.70-fold increase, normalized median fluorescence intensity (MFI) was largely insertion site dependent; the remaining 147G and 156G strains showed approximate increases of 1.54-fold and 1.46-fold, respectively, relative to wild-type (Figure 3A). Strikingly, Bartlett’s Tests revealed that MFI variance was highly strain-dependent (p = 0.71, p = 0.48, and p < 0.0001 for log-transformed 147G, 156G, and all strains, respectively). Given this heteroscedasticity, we analyzed this data using a hierarchical Bayesian model, with variances pooled by insertion site. (Supplementary Figure 13). Based on this model, all engineered strains showed significantly increased MFI compared to wild-type *Synechocystis* (pseudo-p < 0.05 in all cases). These results strongly suggest that SpyCatcher003 display improves SpyTag003-sfGFP binding to *Synechocystis* cells in a largely insertion site-dependent manner.

**Figure 3.**
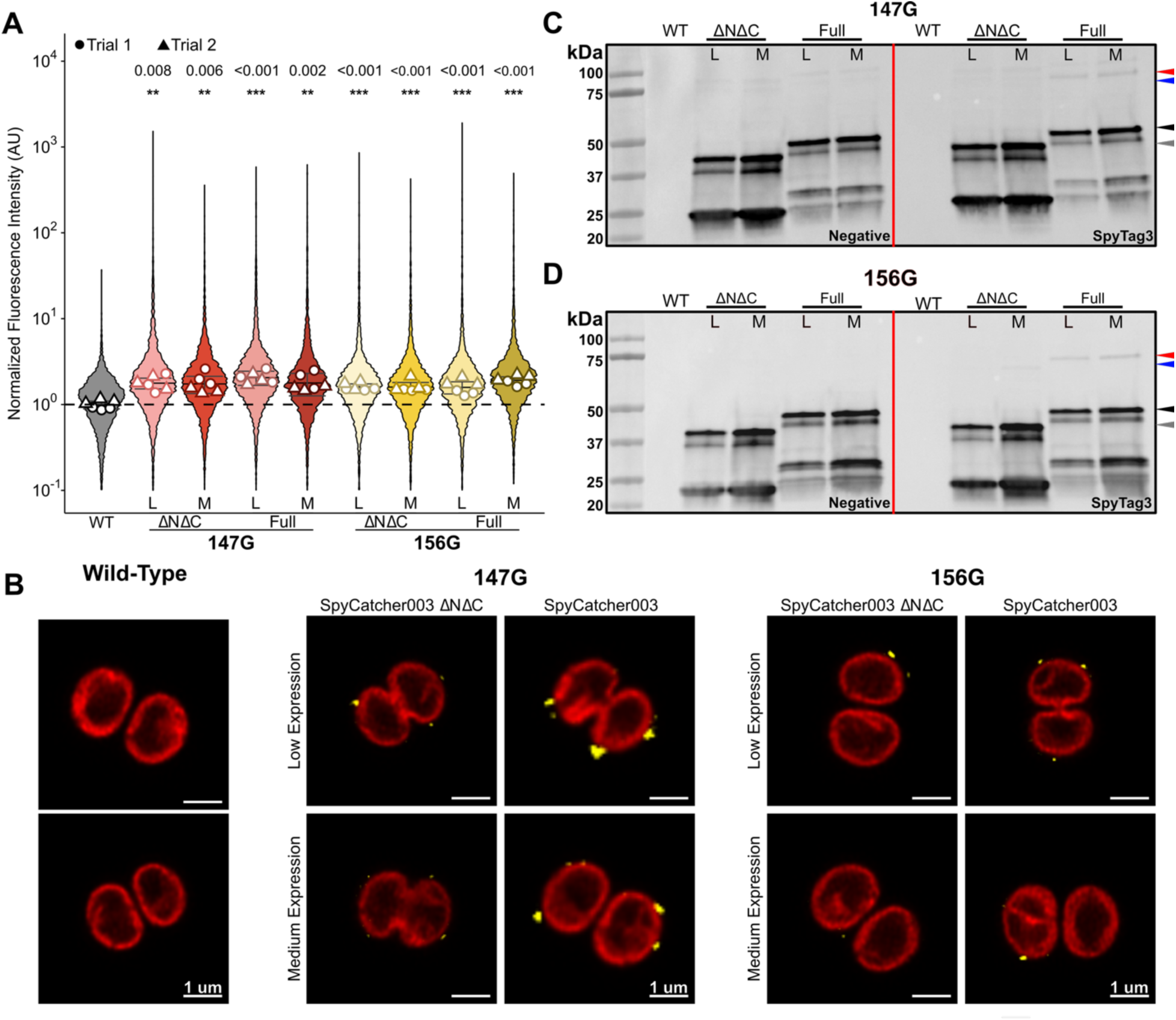
*Synechocystis* strains displaying SpyCatcher003 demonstrate insertion-site dependent reactivity. **A)** Normalized fluorescence intensity of wild-type and PilA1-SpyCatcher003 displaying *Synechocystis* cells, after incubation with SpyTag003-sfGFP as measured by flow cytometry, showing binding of SpyTag003 to the engineered PilA1 strains. Circular and triangular points represent the median fluorescence of individual samples in Days 1 and 2, respectively. Error bars represent the 95% Bayesian Credible interval. Pseudo-p values of median fluorescence compared to wild-type was determined through hierarchical Bayesian Estimation (n =6 biological replicates of ∼100000 cells; *, p < 0.05; **, p <0.01; ***, p<0.001). **B)** Confocal microscopy images of either wild-type (left), PilA1-147G:SpyCatcher003 (center) or PilA1-156G: SpyCatcher003 (right) cells after incubation with SpyTag003-sfGFP (visualized as yellow), showing sfGFP bound to the cell surface. Scale bars represent 1 µm. **C,D)** Anti-SpyCatcher Western blot of WT, PilA1-SpyCatcher003 displaying cells at sites 147G **(C),** and 156G **(D)** after incubation in the absence (left) or presence (right) of SpyTag003-sfGFP (right). Blue and red arrows show SpyTag003-sfGFP covalently bound to PilA1-SpyCatcher003 and PilA1-SpyCatcher003ΔNΔC, respectively. Grey and black arrows represent PilA1-SpyCatcher003 and PilA1-SpyCatcher003ΔNΔC, respectively. Wild-type (WT) cells were included as a negative binding control.

To confirm this binding was localized to the cell surface, we assessed the spatial distribution of the SpyTag003-sfGFP signal relative to *Synechocystis* membrane autofluorescence using confocal microscopy (Figure 3B, Supplementary Figure 14). While wild-type cells exhibited no significant sfGFP fluorescence, all engineered strains exhibited distinct puncta of sfGFP fluorescence close to the cell surface. These puncta were more prominent in strains displaying SpyCatcher003 at insertion site 147G compared to 156G, mirroring the trends seen via flow cytometry. These results confirm site-dependent reactivity of TFP-displayed SpyCatcher003.

Finally, to verify that the increased fluorescence was due to covalent binding, whole cell lysates of the *Synechocystis* strains post-SpyTag003-sfGFP incubation were analyzed by anti-SpyCatcher western blot (Figure 3 C,D). Unbound PilA1-SpyCatcher003 was observed across all engineered samples. When compared to cell-only controls (Figure 3C-D, left), cells incubated with SpyTag003 showed an additional higher molecular weight band at 75 kDa (Figure 3C-D, right). This band is consistent with covalent binding between the PilA1-SpyCatcher003 (44 kDa) and SpyTag003-sfGFP-His-Tag (30 kDa). None of the SpyCatcher003ΔNΔC-displaying strains showed a clear bound band, except for M-156G:SC3ΔNΔC, demonstrating the reduced reactivity of displayed SpyCatcher003ΔNΔC. Anti-His western blot corroborated these results, only showing covalent binding in the strains displaying full-length SpyCatcher003 (Supplementary Figure 15). Taken together, these results strongly suggest that surface-displayed SpyCatcher003 maintained reactivity to SpyTag003 when embedded within TFP. Likewise, strain L-147G:SC3 demonstrated the best reactivity to SpyTag003 across these experiments while maintaining high levels of PilA1 assembly, and was selected for testing within living materials.

### Type IV pilus-mediated display of SpyCatcher003 increases *Synechocystis* binding into the BUD-ELM structure

Type IV Pili are key tools for modulating interactions between bacteria and external cells or materials. To demonstrate this utility, we evaluated our best performing strain L-147G:SC3, for its ability to incorporate *Synechocystis* cells into a macroscopic material formed by the bacterium *Caulobacter crescentus*.^16^ These engineered *C. crescentus* cells, termed a <u>B</u>ottom-<u>Up</u> *<u>D</u>e novo* <u>E</u>ngineered <u>L</u>iving <u>M</u>aterial (BUD-ELM),^16,47^ secrete and display a modified surface-layer (S-layer) protein to form a cell-rich living matrix. To adapt the BUD-ELM for TFP-mediated *Synechocystis* incorporation via covalent SpyTag/SpyCatcher interactions, we first replaced the SpyTag001 peptide displayed in the original matrix with the higher affinity SpyTag003. Additionally, to optimize co-culture with the kanamycin-resistant *Synechocystis* strains and minimize cross-species toxicity, we replaced the *C. crescentus* genomic CdZ operon, which encodes a contact-dependent surface toxin, with a constitutive kanamycin resistance cassette.^48^ Lastly, to facilitate material visualization and analysis, we integrated a constitutive mTurqoise2 expression cassette adjacent to the kanamycin selection marker. These modifications yielded a visually identical BUD-ELM material exhibiting low levels of blue fluorescence (Supplementary Figure 16).

To test how the TFP modification influenced *Synechocystis* incorporation into the BUD-ELM structure, wild-type and L-147G:SC3 *Synechocystis* cells were separately co-cultured with pre-formed BUD-ELM material. To do so, we first established a two-step protocol in which *Synechocystis* and *C. crescentus* were grown separately until exponential phase and BUD-ELM formation, respectively (Figure 4A). Then, *Synechocystis* cells and BUD-ELM material were washed and incubated together in a modified BG11 media (2x BG11, 25 mM HEPES, 12.5 mM Sucrose, pH 7.5), which we formulated to support the growth of both species (Supplementary Figure 17, Supplementary Table 7). This approach enabled the loading of *Synechocystis* cells directly into the BUD-ELM matrix.

**Figure 4:**
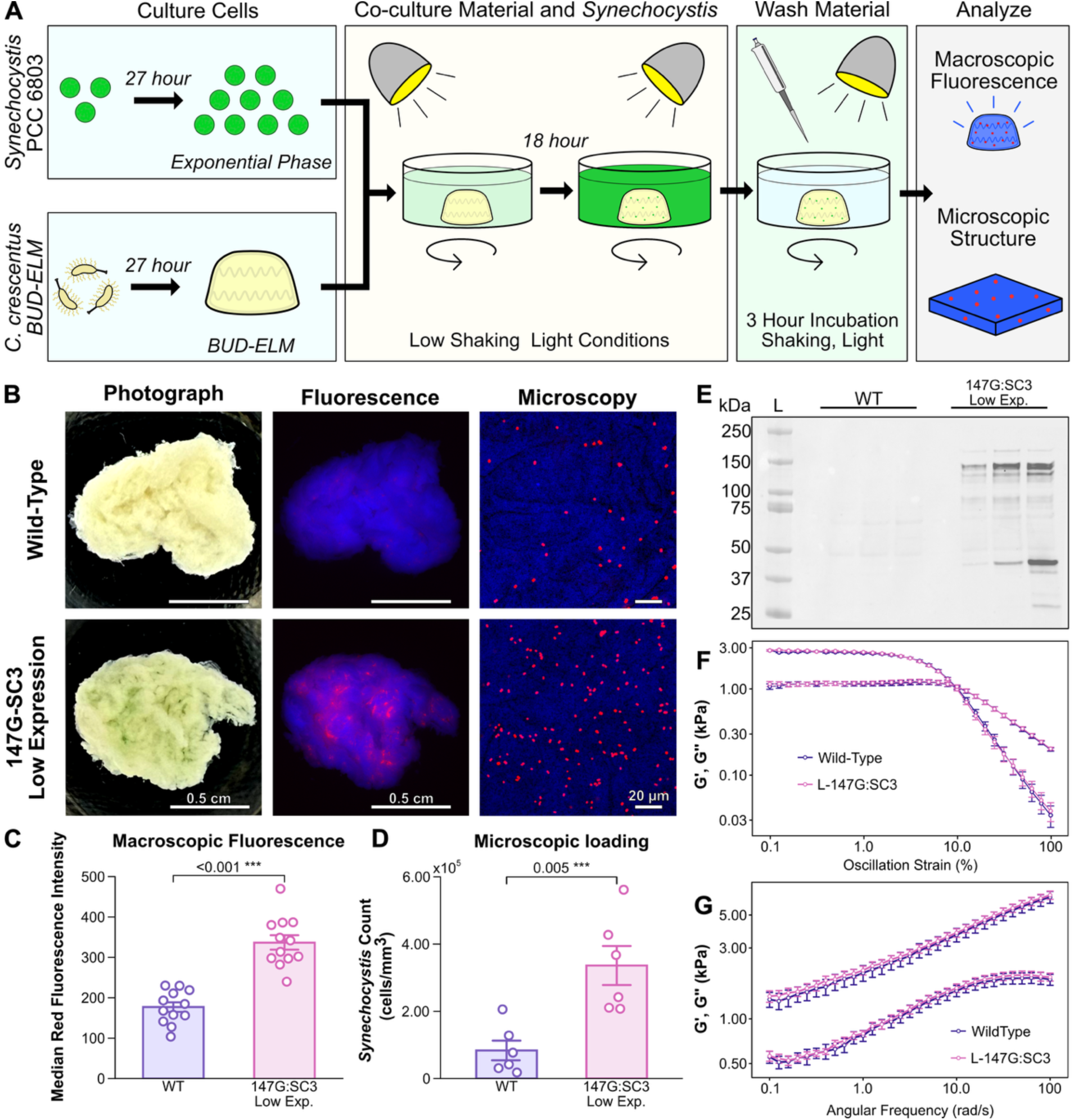
PilA1 display of SpyCatcher003 on the Synechocystis surface improves loading into the BUD-ELM structure. **A**) Schematic illustrating the co-culture of *Synechocystis* and *C. crescentus* materials. *Synechocystis* and *C. crescentus* cells were first grown separately, until exponential phase and BUD-ELM assembly were reached, respectively. Then the cells and material were washed in CC medium and co-cultured for 18 hours under light and shaking conditions, enabling *Synechocystis* to integrate into the BUD-ELM structure. Materials were then washed and imaged to identify *Synechocystis* and *C. crescentus* cells. **B)** Photograph (left), fluorescence macrograph (center), and confocal micrograph (right) images of BUD-ELMs materials after incubation with either wild-type (top), low expression PilA1-147G:SpyCatcher003 displaying (center) or medium expression PilA1-SpyCatcher003 (bottom) displaying *Synechocystis* cells, showing cyanobacterial loading into the BUD-ELM structure. Scale bars for photographs and fluorescence images represent 0.5 cm. Scale bars for microscope images represent 20 µm. **C)** Median red fluorescence intensity of BUD-ELMs after co-culturing with *Synechocystis*, demonstrating that low levels of PilA1-SpyCatcher003 expression improves *Synechocystis* loading. Three biological replicates were measured. **D)** *Synechocystis* loading per cubic millimeter of BUD-ELM material, as determined through Z-stack confocal microscopy, showing that low levels of PilA1-SpyCatcher003 expression improves *Synechocystis* loading, in agreement with the macroscopic fluorescence. Each data point is the median value from nine fields of view across different sections of the material. Three biological replicates were measured. Significance was determined via an unpaired two-sample t-test with a Welch correction for unequal variances. **E)** Anti-SpyCatcher Western Blot of BUD-ELM material after co-culture with *Synechocystis* cells, showing covalent binding between PilA1-147G:SC3 and the material matrix (n=3). **F), G)** Amplitude and Frequency sweeps showing that BUD-ELMs containing Engineered *Synechocystis* maintain their rheological properties compared to wild-type *Synechocystis cells* (n = 6).

After washing the material to remove non-specifically bound cells, we probed *Synechocystis* loading via optical imaging, fluorescence macroscopy, and 3D-confocal microscopy (Figure 4B). The BUD-ELM incubated with the wild-type *Synechocystis* cells was a uniform pale beige, consistent with prior reports.^16^ By contrast, material incubated with L-147G:SC3 developed multiple distinct green patches spanning several millimeters (Figure 4B – left). Fluorescence macroscopy corroborated these findings; wild-type-incubated BUD-ELM possessed low levels of red fluorescence from *Synechocystis* chlorophyll. The L-147G:SC3-incubated materials displayed distinct red puncta, suggesting a high local *Synechocystis* density (Figure 4B – middle). Image analysis revealed a significant increase in mean red fluorescence in the L-147G:SC3 materials compared to wild-type, strongly suggesting enhanced *Synechocystis* loading.

Having observed an increase in *Synechocystis* pigmentation at the macroscopic level, we characterized the density and distribution of *Synechocystis* within the material’s microstructure. The co-cultured BUD-ELMs were gently dissected into ∼0.25 cm^3^ pieces and imaged via Z-stack 3D-confocal microscopy (Figure 4B – right). For each material, we quantified *Synechocystis* density (in cells/ mm^3^) in nine 829 µm x 829 µm x 30 µm fields of view (Figure 4B – right, Figure 4D). *Synechocystis* loading was observed in both wild-type and L-147G:SC3 co-cultured BUD-ELMs, with L-147G:SC3 display improving *Synechocystis* loading by approximately 4-fold (Figure 4C).

Additionally, covalent binding between L-147G:SC3 PilA1 and the BUD-ELM matrix was confirmed via anti-SpyCatcher Western blot (Figure 4E). Replicate material samples exhibited varying intensities of both monomeric and covalently crosslinked L-147G:SC3, consistent with the significant heterogeneity in *Synechocystis* loading observed during confocal imaging. Spatial distribution analysis confirmed non-random cellular clustering, although clustering did not differ significantly between wild-type and engineered strains (Supplementary Figure 18). We speculated that the observed loading heterogeneity was a result of variation in the adhesion affinity and SpyTag003 accessibility across the BUD-ELM structure. These results demonstrate TFP-mediated SpyCatcher003 display as a viable strategy for covalently loading *Synechocystis* cells into material structures.

Lastly, cyanobacteria such as *Synechocystis* PCC 6803 are known to secrete a variety of degradatory enzymes and proteases that may compromise the integrity of the BUD-ELM matrix. To confirm that the L-147G:SC3 mediated increase in *Synechocystis* loading does not degrade the BUD-ELM structure, we measured the mechanical properties of our co-cultured materials by rheology (Figure 4 F,G). Strain amplitude and frequency sweeps of wild-type *Synechocystis* materials were consistent with previous reports of BUD-ELM mechanical properties.^16^ Likewise, L-147G:SC3 BUD-ELM also showed no significant differences in rheological properties compared to wild-type materials or reported BUD-ELM values. Taken together, these results demonstrate that TFP-mediated display can increase incorporation of *Synechocystis* cells into living structures without compromising the material’s rheological properties.

## Discussion

Type IV Pili show tremendous potential as a widely distributed platform for bacterial surface display, but the ability to display proteins without impairing TFP assembly has remained an open challenge. To bridge this gap, we have demonstrated a computational strategy for domain insertions into Type IV Pili, designed to preserve TFP assembly across both monomeric and multimeric structures. Using the major Type IV pilin PilA1 of the cyanobacterium *Synechocystis* PCC 6803 as a test case, we predicted and validated the insertion tolerance of SpyCatcher003 across multiple sites. Consequently, we found insertion site, cargo size, and fusion pilin expression level as critical factors for proper display, while confirming that PilA1-fused SpyCatcher003 retained covalent reactivity on the *Synechocystis* surface. Lastly, we showed that TFP-mediated SpyCatcher003 display significantly enhances *Synechocystis* loading into a living material structure via covalent SpyTag/SpyCatcher associations, demonstrating our system’s utility in expanding the role of TFP for user-defined applications.

Importantly, centering our strategy around maintaining TFP assembly allowed us to fully leverage the structural homologies across TFP. This design choice enables accurate prediction of insertion tolerant regions despite the small available dataset, which limits current machine learning-based methods. Likewise, by focusing our analysis on direct structural information, our framework is agnostic to specific protein structure prediction tools and can readily integrate future tools that accurately predict multimeric TFP assemblies. These considerations also make our strategy chassis-independent, and generalizable across diverse native TFP systems.

Our approach has also yielded multiple improvements in TFP display over the current state of the art. Compared to previous attempts utilizing C-terminal fusion of a small 6.5 kDa affibody to *Synechocystis* PilA1, our best performing sites (147G and 156G) maintained 6-8 times more cells in suspension after 16 hours, despite displaying a 92.8% larger cargo protein.^23^ These improvements signal that optimized insertion into PilA1 better preserves the structural integrity of the TFP fiber, while significantly expanding display capacity. Leveraging this expanded display capacity also allowed us to augment TFP-mediated cellular interactions with the extracellular environment. Our optimized SpyCatcher003-displaying strains enabled covalent binding to both soluble probes and cell-based materials, expanding the toolkit for *Synechocystis* display and patterning.

These gains were mirrored when binding *Synechocystis* cells to surfaces. Compared to the binding of gold nanoparticles via C-terminal peptide display, which required a hyper-pilated, TFP retraction-negative *Synechocystis* mutant for significant binding,^24^ internal SpyCatcher003 display alone was sufficient for improved *Synechocystis* loading into a living material structure. As such, we believe that optimized TFP display can enable broad functionality with minimal changes to native TFP machinery, which will also improve generalization of this method across chassis.

Despite these key enhancements to TFP display, our approach possesses both limitations and opportunities for improvement. Within this study we only tested four sites, and although 75% of sites matched our predictions for insertion tolerance, a deeper screening approach would further validate our model. Analyzing the one site that our pipeline incorrectly predicted as insertion tolerant, 150T, we observed that it fell within a local minimum for both solvent accessibility and flexibility. This observation suggests that our criteria or their thresholds are incomplete. Further work to increase screening throughput and fine tune criteria thresholds could optimize our workflow and improve the predictive power of our approach.

There is also potential for further optimization of the PilA1 display system. Our workflow did not factor in either fusion pilin expression level or cargo size, which our experimental work confirmed to be critical parameters for display. These limitations could be ameliorated by tailoring our pipeline to account for varying expression levels of engineered pilin within the TFP fiber, while accounting for TFP stability as fusion pilin levels increase. Likewise, we observed a low abundance and rate of glycosylation of engineered pilins across all tested constructs, potentially suggesting protein instability and unknown regulation of TFP production. These issues may have also been exacerbated by our use of a non-motile strain of *Synechocystis*, and more work must be done to understand how our engineering approaches may impact motile strains that may possess altered TFP. In tandem, improving our understanding of TFP regulation, such as potential pilin proteases or stabilizing post-translational modifications, will enable improvements to TFP display capacity while minimizing burdens to cell fitness.

Ultimately, this work begins to elucidate design rules for stable display on TFP and builds a framework for validating TFP display systems. By applying this framework to *Synechocystis*, we successfully stabilized the TFP-mediated display of SpyCatcher003, enabling covalent crosslinking between *Synechocystis* and exogenous proteins and materials. Moving forward, TFP represent highly promising targets for near-universal protein display, offering a pathway towards programmable actuation across the bacterial kingdom.

## Materials and Methods

### Bacterial growth conditions

*Synechocystis* PCC 6803 (Substrain GT-S) cells, unless otherwise noted, were cultivated in BG11 medium (Sigma-Aldrich) buffered to pH 7.8 with 25 mM HEPES (Sigma-Aldrich), supplemented with 50 μg/ml kanamycin (Sigma-Aldrich) when appropriate. For growth on solid media, individual colonies from cell culture were first streaked onto BG11 plates containing 1.5% (wt/vol) Bacto agar (BD) and 3 g/L sodium thiosulfate (Sigma-Aldrich). BG11 plates were incubated in Minitron shaking incubators at 28 °C under ∼50 μmol m⁻² s⁻¹ illumination, 1.6% (vol/vol) CO_2_, 55% relative humidity, under static conditions. To minimize evaporation, BG11 plates were sealed with Parafilm during cultivation, and after cells reached sufficient density, plates were maintained under low illumination at room temperature for up to 2 weeks.

To prepare *Synechocystis* cells for experiments, individual colonies were first streaked onto BG11 plates, and grown for ∼48 hr into large colonies ∼2.5 mm in diameter. These large colonies were then inoculated into 1.5 mL BG11 medium in 12 well culture plates. Cultures were cultivated in Minitron shaking incubators at 28 °C under ∼50 μmol m⁻² s⁻¹ illumination, 1.6% (vol/vol) CO_2_, 55% relative humidity, and 190 rpm shaking with a 25 mm orbit, and grown to an OD_730_ =∼5. To minimize the impacts of evaporation, ∼300 µL of fresh BG11 medium was added to these cultures after ∼36 hours of growth. After approximately 60 hours, cells were pelleted via centrifugation (1000 rcf, 3.5 min), resuspended into fresh BG11, and diluted to a final OD_730_=0.15 in 25 mL or 50 mL sterile Erlenmeyer flasks. Cultures were then grown at 28 °C under ∼50 μmol m⁻² s⁻¹ illumination, 1.6% (vol/vol) CO2, and 190 rpm shaking with a 25 mm orbit until a desired density was reached.

*Escherichia coli* DH10B (New England Biolabs) was cultivated in LB media (Sigma-Aldrich) for liquid growth or LB media with 1.5% (wt/vol) agar (Fisher Bioreagents) for growth on solid media. When antibiotics were needed, media was supplemented with 50 μg/mL kanamycin.

*C. crescentus* BUD-ELM cells, unless otherwise noted, were cultivated from individual colonies in 32 mL of PYE medium in 125 mL Erlenmeyer flasks. Flasks were incubated in New Brunswick Innova 44 shaking incubators with a 2-inch orbit, shaking at 250 rpm at 30 °C. For growth on solid media, individual colonies were streaked onto PYE agar plates and incubated at 30 °C.

### Plasmid Assembly and Transformation

All plasmids and genetic parts used in this work are listed in Supplementary Tables 2 and 3. The RSF1010 origin of replication, pNPTS138 *C. crescentus* integration backbone, kanamycin resistance, spectinomycin resistance, and terminators were stored as individual plasmids containing BsaI cut sites. Genetic sequences for the s6 and s7 promoters, PilA1, SpyCatcher, ELP_60_, SpyTag003, and mTurquoise2 were obtained via Twist Biosciences, and were flanked by BsaI cut sites. Plasmids for genomic integration in *C. crescentus* also included 800 bp homologous recombination regions flanking each end of the insertion sequences of interest, corresponding to the RsaA gene (CCNA_01059) and CdZABCDI operon (CCNA_00718-CCNA_00721) for BUD-ELM and KanR-mTurquoise2 integration, respectively.

Plasmids were constructed by Golden Gate Assembly and transformed into *E. coli* DH10B via heat shock. Transformed cells were grown on LB-kanamycin agar plates, and individual colonies were grown in LB-kanamycin cultures overnight. Plasmid DNA was purified from *E. coli* cells via miniprep extraction (New England Biolabs), and sequence verified via nanopore sequencing (Plasmidsaurus).

The transformation procedure for *Synechocystis* was adapted from Cengic et al. 2018.^23^ Cells from an independent colony were first grown (10 mL per transformation) cultures to exponential phase (OD730 = 0.6-08). Exponential phase cells were placed on ice, washed 3 times in sterile cold water by centrifugation (10 minutes, 4000 rpm, 4 °C) concentrated to 1% of the original culture volume in cold water, and separated into 50 μL aliquots on ice. For each transformation, ∼350 ng of plasmid DNA was added to an aliquot, up to a maximum DNA volume of 5 μL, and the mixture was incubated on ice for 10 minutes. Aliquots were then transferred into cold electroporation cuvettes (2 mm gap, Bio-Rad) and electroporated using a Bio-Rad Micropulser on the EC2 setting (2.5 KV). Immediately following electroporation, 500 μL BG11 was added to each cuvette. This mixture was then spread onto a sterile 0.45 µm nitrocellulose membrane (Sigma-Aldrich) (placed on top of a BG11 agar plate and incubated in light conditions for 24 hours. These membranes were then transferred to BG11-kanamycin plates, which were sealed with parafilm, and then incubated in light conditions. Transformed colonies were visible after ∼7-10 days.

### Transformation and Genomic Integration into the *C. crescentus* genome

To prepare *C. crescentus* cells for electroporation, individual colonies of *C. crescentus* BUD-ELM cells were first inoculated into 30 mL of PYE medium in 135 mL Erlenmeyer flasks and grown at 30 °C under shaking conditions for 24 hours, until stationary phase was reached. Then, 3 mL of cells were pelleted at 4°C, washed 3 times in cold, sterile, ultrapure water, and resuspended into 50 µL of cold, sterile, ultrapure water. Washed cells were stored on ice between steps. Next, ∼100 ng of plasmid (totaling less than 5 µL in volume) was gently added to the washed cells, and the mixture was transferred to a 1 mm electroporation cuvette (Bio-Rad). Cells were then transformed via electroporation using a BioRad Micropulser with a 12 kV/cm field strength. Immediately following electroporation, 1 mL of fresh PYE medium was added to the cuvette, and the mixture was transferred to a sterile 1.5 mL tube, which was then incubated at 30°C under shaking conditions for 4 hours. After incubation, the cell mixture was then spread on PYE agar plates containing 25 µg/mL kanamycin (Sigma-Aldrich); these plates were incubated at 30 °C under static conditions until colonies were visible (∼3-5 days). From the transformation plate, individual colonies were then inoculated into 30 mL of PYE medium in 135 mL Erlenmeyer flasks and grown at 30 °C under shaking conditions for 24 hours, until stationary phase was reached. Cells were then plated on PYE supplemented with 3% w/v sucrose to select for excision of the plasmid and sacB gene, leaving the target sequence in the genome. Integration of the sequences was confirmed by colony PCR using a touchdown thermocycling protocol with an annealing temperature ranging from 72–62 °C, decreasing 1 °C per cycle. The PCR amplicons have been fully sequenced.

### Multiple sequence alignment of pilins

Multiple Sequence alignment was performed between the reference TFP sequences and the *Synechocystis* PCC 6803 PilA1 pilin via Clustal Omega.^49^ Sequences were visualized using Jalview.

### Generation of the *Synechocystis* PilA1 monomer

The PilA1 monomeric structure was predicted via the Alphafold3 webserver.^27^ To better compare this monomer against known TFP structures, the known N-terminal secretion signal was removed from the PilA1 sequence prior to structure generation.

### Prediction of Interface residues in the *Synechocystis* PilA1 monomer

To analyze the interfaces of known TFP structures, multimeric structures were first acquired from the Protein Data Bank. Inter-chain interactions for each structure were calculated by constructing a k-d tree and executing a K-nearest neighbor search from one chain to each other chain, with a radius of 4.5 Å. This process yielded a unique set of interface residues for each structure. In parallel, a monomer from the center of the multimeric fiber was extracted and truncated to remove the N-terminal ∼20 residues up to and including the highly conserved proline, which signifies the boundary between segments of the N-terminal helix domain. This step was taken to avoid misalignments to the *Synechocystis* PilA1 monomer, as this N-terminal segment frequently becomes elongated in multimeric TFP compared to isolated monomers.

Next, the generated PilA1 structure was aligned against each truncated reference monomer by Template Monomer alignment (TM-align), and the resulting TM-scores, normalized to the length of the reference structure, were stored in a TM-vector 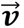. To generate a set of projection interface scores, The alignment was then analyzed residue-wise in the following ways:

1. Alignment was scored on [0, 0.5, 1], representing a gap, partial alignment, and full alignment, respectively. The resulting scores were stored in an alignment matrix.
2. Aligned residues from the reference structure were cross referenced against the corresponding interface residue list, and scored on [0, 1] for non-interface and interface residues, respectively. The resulting scores were stored in an interface matrix.

These operations yielded two n x m matrices, an alignment matrix ***A_prj_*** and a contact matrix ***C***, where n is the number of residues in the PilA1 sequence, and m is the number of aligned reference structures. Additionally, a normalization vector 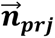 was created by calculating the ratio of contact to aligned residues for each position. To generate a final interface prediction score, the following operations were performed:

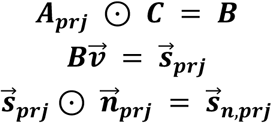

Where ***A_prj_*** is the alignment matrix, **C** is the contact matrix, ***B*** is the aligned contact matrix, 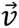 is the TM-vector, 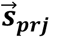 is the interface scoring vector, 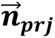 is the normalization vector, and 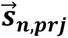 is the normalized scoring vector. These resulting interface prediction scores were then projected onto the PilA1 monomer to depict predicted interface residues on the structure.

Likewise, to generate the set of insertion interface scores, a central chain within each TFP multimer was first replaced with a trimmed PilA1 monomer. PilA1 interfaces for each inserted monomer were then directly calculated via a K-nearest neighbor search with an interatom distance of 4.5 Å, yielding matrix **A_ins_**. These measurements were then combined by a weighted average using their respective TM- scores (normalized to the length of PilA1) as weights, normalized by the total sum of TM-scores to create a second predicted set of interface residues:

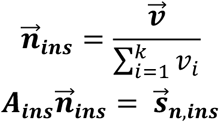

Where 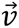 is the TM-vector, 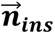 is the insertion normalization vector, and 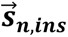 is the set of insertion interface scores.Residues that received a score greater than 0.2 in both prediction methods were considered to be interface residues and excluded from the list of viable insertion sites.

### Measurement of Solvent Accessible Surface Area (SASA)

To calculate the solvent accessible surface area (SASA) of each PilA1 residue, monomers previously inserted into the TFP reference structures were analyzed using the Shrake-Rupley algorithm.^36^ SASA values for each residue were then weighted across measurements by TM-score and normalized by total sum of TM-scores 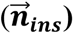. Lastly, a sliding average was calculated using the two residues before and after each site. Sites with an above average SASA score (0.200) relative to the whole monomer were considered to be solvent accessible for downstream analysis.

### Measurement of Residue Root Mean Squared Flexibility (RMSF) using CABS-Flex

Root mean square flexibility (RMSF) of PilA1 monomers inserted as a central chain into each reference TFP multimer were calculated using CABS-Flex under the Rigid-pLDDT Flexibility mode.^37^ To ensure that flexibility was measured in the proper context, PilA1 monomers that were previously inserted into reference TFP structures were used for all calculations. To consider local flexibility effects, a sliding window was created such that the value at each residue represents the average RMSF of the two preceding and three subsequent residues, for a total window size of 6. These measurements were performed with all 8 reference TFP structures, and TM-scores were used to calculate a weighted average for each residue 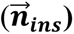. All residues with a RMSF above the mean (0.588 Å) were considered flexible for insertion.

### Selection of PilA1 insertion sites

To create a final set of insertion sites for experimental validation, the binarized interface SASA, and RMSF values were combined to produce a set of non-interfacial residues with above average surface accessibility and flexibility. Regions with less than 3 contiguous viable sites were excluded, as it was determined that insertions might disrupt the surrounding area. Then, the largest contiguous region was considered as the optimal site for insertion, namely 144A-157G. Three sites were chosen from this region for downstream testing.

### Sedimentation Assay of *Synechocystis* cells

*Synechocystis* cells inoculated into 10 mL cultures at an initial OD_730_ = 0.15 and cultivated for ∼48 hours, until OD_730_ reached ∼2. Cells were pelleted and resuspended into fresh BG11 media. Cells were then diluted to 5 mL with BG11 into glass tubes, to a final density OD_730_ = 2. Sedimentation was monitored at specified time points by sampling 150 μL from the topmost layer and measuring OD_730_ in 96 well plates via a Tecan Spark plate reader.

### Growth curve analysis of *Synechocystis* cultures

*Synechocystis* strains grown from independent colonies into 12-well tissue culture plates were washed and diluted to OD_730_= 0.15 in 10 mL BG11 in 50 mL Erlenmeyer flasks. Flasks were then placed in the shaking incubator under standard light conditions. At specified time points, flasks were removed from the shaker, 200 µL aliquots transferred to 96 well plates, and OD_730_ was measured using a Tecan Spark Microplate reader.

### SpyTag-sfGFP binding to *Synechocystis* cells

*Synechocystis* strains grown from independent colonies into 12-well tissue culture plates were washed, diluted to OD_730_= 0.15 in 10 mL BG11 in 50 mL Erlenmeyer flasks, and cultivated until OD_730_ = ∼1.2 (approximately 36 hours into growth). Cells were then pelleted, washed once in phosphate buffered saline (Fisher Bioreagents) containing 0.1% v/v Pluronic F-108 (Sigma-Aldrich), dubbed PBS-P, and resuspended in 750 µL PBS-P at OD_730_ = 1 containing 200 nM SpyTag003-sfGFP. Samples were then incubated for 6 hours at room temperature in low-light conditions with gentle rotating before being pelleted, washed twice, and resuspended in PBS-P.

### Extraction of Type IV Pili from *Synechocystis* cell surface

To prepare S*ynechocystis* cells for TFP extraction, 50 mL of culture were grown to OD_730_ = ∼2.84. Cells were then diluted to 50 mL in conical tubes, such that the cells in all tubes contained the same OD730 value to that of the lowest density culture. These cells were pelleted, resuspended in 0.5 mL of 0.5x BG11 medium, transferred to 1.5 mL tubes, and vortexed at max speed for 5 minutes. To remove the cells, samples were then centrifuged for 5 minutes at 14000x g, and the supernatant was transferred to new tubes. This process was repeated a total of 3 times to ensure that all cell mass was removed. Proteins were precipitated via tricholoracetic acid (Thermo Scientific), which was added to the supernatant to a final 10% (w/vol) before incubating for 2 hours on ice. Samples were then centrifuged for 10 minutes at 14000x g, and the resulting pellet was washed 2 times with cold 90% acetone. After the second wash, the open tubes were incubated in sterile conditions for 15 minutes to evaporate excess acetone and stored at -20 °C until analysis.

### SDS-PAGE analysis of extracted *Synechocystis* pili

To characterize the extracted TFP, extraction pellets were resuspended into 50 µL of 2x Laemmli buffer (BioRad) containing 1x of β-mercaptoethanol (Sigma-Aldrich), which were incubated at 95 °C for 5 minutes before being diluted 1:1 with 1x PBS, to a total volume of 100 µL. Then, 25 µL of each sample was loaded onto a Criterion TGX Stain-Free™ gel (Bio-Rad) and analyzed via SDS-PAGE. After running, the gel was incubated in ultrapure water at room temperature and allowed to incubate under mild shaking for 5 minutes. This process was repeated a total of three times. Then, to visualize the protein bands, the gel was incubated with GelCode Blue Stain reagent (Thermo Fisher) at room temperature under mild shaking for 1 hour. After incubation, the stain was replaced with ultrapure water, and the gel was incubated for an additional hour. The washed protein gel was then imaged using a BioRad GelDoc Go imager.

### Western Blot Analysis

For Western blot analysis, proteins resolved via SDS-PAGE were transferred onto 0.2 μm nitrocellulose membrane (Bio-Rad) and blocked for 1 h at room temperature with SuperBlock™ blocking buffer (Thermo Scientific). Membranes were then washed four times in Tris-buffered Saline (Sigma-Aldrich) with 0.1% v/v Tween 20 (Sigma-Aldrich) (TBST) before incubation in antibody-TBST solution containing 1:5000 dilution of either Monoclonal rabbit Anti-SpyCatcher® antibody (Bio-Rad) or Anti-HisTag-HRP Antibody (Thermo-Fisher) for 45 minutes at room temperature. Membranes were washed four more times with TBST, and if Anti-SpyCatcher was used, were incubated in a 1:500- dilution of Monoclonal goat anti-rabbit-HRP secondary antibody (Bio-Rad) for 45 minutes. Four final TBST washes were performed, and chemiluminescence was activated by applying Clarity Max Western ECL Substrate (Bio-Rad). Chemiluminescence and Visual images were captured using a Bio-Rad GelDoc Go system. For whole-cell analysis, samples were first normalized by cell density (OD_730_) before lysis in Laemmli buffer at 98 °C for 10 minutes. For analysis of the BUD-ELM material, approximately 5 µL segments of material were incubated in 50 mL of Laemmli buffer (Bio-Rad), incubated at 98 °C for 10 minutes, and gently centrifuged to separate remaining solids. 25 µL of each sample was loaded for analysis.

### Expression of SpyTag003-sfGFP

pET28a::SpyTag003-sfGFP with a C-terminal hexa-histidine (His_6_) tag was transformed into chemically competent *E. coli* BL21 (DE3) cells and plated on LB agar (Sigma-Aldrich) supplemented with 50 ug/mL kanamycin (Sigma-Aldrich). Single colonies were picked and inoculated into 15 mL Terrific Broth (Sigma-Aldrich) medium supplemented with 50 µg/mL kanamycin, incubated at 37 °C with shaking at 200 rpm for 16 hours. The culture was then inoculated into 500 mL cultures of Terrific Broth supplemented with 50 µg/mL kanamycin, incubated at 37 °C with shaking at 200 rpm until OD_600_ 0.5–0.8, when the cultures were induced with 0.5 mM isopropyl β-D-1-thiogalactopyranoside (IPTG) (GoldBio) to trigger protein expression. Induced culture was grown for 18 h with shaking at 180 rpm at 18 °C. Cells were harvested by centrifugation at 8000×*g* for 15 min at 4 °C prior to purification.

### Purification of SpyTag003-sfGFP

Approximately 16 g of frozen cell pellet was thawed in 80 mL of binding buffer (50 mM Tris-HCl pH 8 (Sigma-Aldrich), 30 mM imidazole (Sigma-Aldrich), 400 mM NaCl (Sigma-Aldrich), 5% glycerol (Sigma-Aldrich), 1 mM dithiothreitol (DTT) (GoldBio)). Resuspended cells were incubated 100 ug/mL Lysozyme (Sigma-Aldrich), DNase I (Sigma-Aldrich), and RNase (Sigma-Aldrich) at 4°C with stirring until homogenous. The lysate was then rotated at room temperature for 30 min and mechanically disrupted using a Qsonica Q500 sonicator. Lysate was subsequently clarified at 70000 x g for 1 hour at 4 °C, and supernatant was loaded onto equilibrated Ni 2+ Sepharose 6 Fast Flow column (Cytiva) for affinity purification.

All purification steps were performed at 4 °C. Bound protein was washed with 10 column volumes of binding buffer to remove residual contaminants and eluted with elution buffer (20 mM Tris-HCl pH 8, 200 mM imidazole, 350 mM NaCl, 1 mM DTT). Sample was concentrated and exchanged into storage buffer (20 mM Tris-HCl pH 7.5, 300 mM NaCl, 5% glycerol) using an Amicon® Ultra Centrifugal Filter, 3 kDa Molecular Weight Cutoff (Sigma-Aldrich). Sample purity was assessed using SDS-PAGE and concentration was determined by the Bradford Assay (Bio-Rad).

### Flow Cytometry analysis of *Synechocystis* cells

The fluorescence of *Synechocystis* cells incubated in sfGFP was measured via Flow Cytometry (Sony SH100S, FL1 channel excitation 488 nm emission 525 nm). A total of 100000 events were collected per sample Cells were gated based on FSC-A by SSC-A plots, using wild-type *Synechocystis* as a reference. Then, these scale values were exported from FlowJo as csv files and analyzed in R. To correct for batch effects between days, the full data for each sample was normalized by the mean of the median fluorescence intensity (MFI) of the wild-type samples collected on that day. Variances within the strains for sites 147G and 156G, as well as the variances for all samples, were first analyzed by Bartlett’s test, which confirmed insertion site dependent variance (p = 0.71, p = 0.48, and p < 0.0001 for log-transformed 147G, 156G, and all strains, respectively).

To evaluate the display efficacy of each strain, MFI values were analyzed using a hierarchical Bayesian heteroscedastic Gaussian model. In this model, Population means (ꞵ) were estimated for each strain, whereas variances (*σ*) were pooled by insertion site (Wild-Type, 147G, 156G). The joint posterior distribution was defined as:

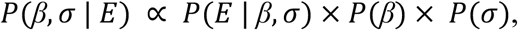

Where E represents the observed experimental data, and the log-transformed MFI data follows a Gaussian likelihood:

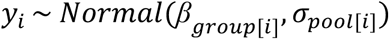

A log-link function was applied to the variance component (ln(σ_pool_) ∼ 0 + σ_pool_) to ensure standard deviation remained positive.

To prevent over-parameterization, we used weakly informative, regularizing prior distributions, transformed into log-space, to bound the parameter landscape. To specify a prior for population group means, we leveraged prior data from C-terminal Z_Taq_ display on PilA1, which reported an approximate ∼2.5 fold increase in MFI. This value was log-transformed and set to represent 2 deviations above the mean (ln(2.5)/2 = 0.458), which was applied to model population group means within a Gaussian prior as *β ∼ Normal*(*0*, *0.458*). For the residual variance parameters, we applied a Gaussian prior as *ln*(*σ*) ∼ *Normal*(–*1, 0.5*), which targeted a standard deviation of approximately 0.36, to prevent singular fits.

Joint Probability was sampled using the Hamiltonian Monte Carlo algorithm. Four independent Markov Chains were simulated for 2000 iterations with a 1000-iteration warm up phase per chain, yielding a total pool of 4000 post-warmup posterior draws for statistical inference. Convergence of all parameters was confirmed via the Gelman-Rubin diagnostic (R-hat = 1.00). Model adequacy and prior calibration were visually audited using Prior and Posterior Predictive Checks to ensure that the posterior simulations accurately captured the underlying density profile of the data. To analyze the directional shifts between the engineered PilA1 groups and the wild-type control, the differences in posterior draws were measured for each of the 4000 HMC draws, and quantified using the probability of direction (pd) metric. This metric was converted into pseudo-p-values (p = 2* (1 - pd)) to better describe conventional alpha significance thresholds.

### Confocal Microscopy of *Synechocystis* cells

*Synechocystis* strains grown from independent colonies into 12-well tissue culture plates were washed, diluted to OD730= 0.15 in 10 mL BG11 in 50 mL Erlenmeyer flasks, and cultivated until OD_730_ = ∼1.2 (approximately 36 hours into growth). Cells were then pelleted, washed once in PBS-P buffer, and resuspended in 750 µL PBS-P at OD730= 1 containing 200 nM SpyTag003-sfGFP. Samples were then incubated for 6 hours at room temperature in low-light conditions with gentle rotating before being pelleted, washed twice in PBS-P, and resuspended in 1 mL PBS-P. 1.5 µL of washed cells was then placed onto a No. 1.5 coverslip and covered with a slab of PBS-Agarose (2% w/v). Cells were then imaged using a Nikon Ti2 Eclipse Super-Resolution Microscope in super-resolution made, using the FITC (1% power) and Cy5 (0.3% power) channels for sequential visualization of sfGFP and chlorophyll fluorescence, respectively. Images were deconvoluted in NIS Elements and analyzed using ImageJ.

### Biolog Carbon Utilization Assay of *C. crescentus growth*

To screen for *C. crescentus* growth in different carbon sources, individual colonies were first inoculated into PYE medium and grown for 24 hours until stationary phase was reached. Then, cells were washed 3 times in 2x BG11 (pH = 7.5) medium, before resuspension to a final density of OD_600_ = 0.001 in 2x BG11 medium. Washed cells were then distributed in 150 µL aliquots into each well of a Biolog PM1 or PM2A Carbon Utilization Assay plate. Each Assay plate was then placed into a humidity cassette and incubated in a Tecan Spark microplate reader at 30 °C under double orbital shaking conditions for 48 hours, during which OD600 measurements were recorded every 20 minutes via a Tecan Spark plate reader. Growth Curves were then analyzed using the GrowthCurver package in R.

### Growth Curves to measure *C. crescentus* response to sucrose

To measure the growth of *C. crescentus* cells, cultures were first individual colonies were first inoculated into PYE medium and grown for 24 hours until stationary phase was reached. Then, cells were washed 3 times in 4x BG11 (pH = 7.5) medium, before resuspension to a final density of OD_600_ = 0.002 in 4x BG11. 75 µL of cells were then distributed into wells of a 96-well microplate and diluted to 150 µL with sucrose solution such that each well contained 0-20 mM sucrose in 2.5 mM increments. Microplates were then placed into a humidity cassette and incubated in a Tecan Spark microplate reader at 30 °C under double orbital shaking conditions for 48 hours, during which OD600 measurements were recorded every 20 minutes. Growth Curves were then analyzed using the GrowthCurver package in R.

### Co-culture between *Synechocystis* and BUD-ELM material

To co-culture *Synechocystis* cells with the BUD-ELM material, *Synechocystis* and *C. crescentus* BUD-mTK cells were cultured independently in BG11 or PYE media for ∼27 hours, until mid-exponential phase (OD_730_ = ∼0.8) or complete BUD-ELM formation, respectively. *Synechocystis* cultures were then harvested, washed twice and resuspended to OD_730_ = 0.8 in fresh CC media via centrifugation. Likewise, once *C. crescentus* cultures produced BUD-ELMs, the material was isolated and was gently washed in fresh CC media 3 times via pipette. Then, the BUD-ELM materials were incubated in 2 mL of the washed *Synechocystis* cells for 18 hours at 30 °C under light conditions (∼50 μmol m⁻² s⁻¹) with increased CO_2_ (1.6% v/v) and mild shaking (85 rpm, 25 mm orbit). After incubation, the *Synechocystis*-containing CC medium was removed, and the BUD-ELM-mTK material was gently washed with fresh CC medium three times before being resuspended in 2 mL of CC medium. To remove weakly-attached cells, the material was then transferred back to the shaking incubator and incubated for an additional 3 hours under identical conditions. After this second incubation the material was once again removed, washed three times with fresh CC medium, and resuspended in CC medium. This material was then analyzed via fluorescence macroscopy, 3D-confocal microscopy, Western Blot and rheology.

### 3D Confocal Macroscopy of BUD-ELM Material

BUD-ELM material was imaged using a Nikon SMZ800 stereo microscope. Red and Blue fluorescence (were captured sequentially with 300 ms and 400 ms exposure, respectively, and a gain of 11.4x. Optical microscopy data were acquired using the software NIS-Elements AR (version 4.51.01). All pictures presented were generated using ImageJ software, and the visualized intensities for each channel were normalized across samples. For analysis of *Synechocystis* loading, the blue fluorescence images were segmented in MATLAB 2025A to create binary masks of the material area. The resulting binary masks were applied to the red fluorescence images, which were then used to calculate the mean red fluorescence intensity, which was subtracted from the surrounding background intensity before analysis.

### Confocal Microscopy of BUD-ELM material

BUD-ELM material co-cultured with *Synechocystis* cells was first washed twice in 1 mL CC medium by gentle pipetting and then resubmerged in 2 mL sterile CC medium. Then, three pieces of each BUD-ELM material, approximately 5 mm in diameter, were gently dissected from the structure. These pieces were placed onto No. 1.5 glass coverslips (Avantor) and sandwiched underneath an agar pad (2% w/v). Images were acquired using a Nikon Ti2 Eclipse Super-Resolution Microscope in super-resolution mode, using DAPI (4% power) and Cy5 (0.3% power) to image *C. crescentus* and *Synechocystis* cells, respectively. For each image, we collected 101 Z-stack images (0.3 µm separation between stacks) of each field of view, resulting in images with a total volume of 829.44 µm x 829.44 µm x 30 µm (0.021 mm^3^). For each piece of material, we imaged three fields of view, with a total of nine fields of view per sample. Data were acquired using the NIS Elements software (AR 6.02.01, 64-bit).

Z-stack images were analyzed using MATLAB 2025A. For each image, we first separated the DAPI and Cy5 Channels to isolate *C. crescentus* and *Synechocystis* cells, respectively. To filter out artefacts, an initial minimum intensity threshold of 2 and 170 were applied, respectively.

To quantify *Synechocystis* loading into the BUD-ELM structure, the number of cells per image was first quantified by image segmentation followed by regionprops3, to provide the list and number of objects present in the sample. To avoid small artefacts that passed the initial threshold, we applied a volume threshold of 20 voxels (∼3.9 µm^3^), which is smaller than the size of a *Synechocystis* cell. The number of remaining cells was then used as a measurement.

To quantify the amount of material imaged, we first calculated the total volume in the DAPI channel using the total sum of voxels above the set threshold, multiplied by the dimensions of each voxel (0.81 µm x 0.81 µm x 0.3 µm) to yield a volume in µm^3^. This value was then converted to mm^3^ for reporting.

To calculate *Synechocystis* loading for each field of view, the calculated number of cells was normalized by the material volume to obtain a measurement in cells/mm^3^. To get a holistic measurement for each piece of material, we calculated the median cells/mm^3^ value for each strain, which was then reported.

### Rheological measurements of co-cultured BUD-ELM material

The rheological properties of BUD-ELMs co-cultured with *Synechocystis* cells were evaluated on a strain-controlled rheometer (TA Instruments ARES-G2) equipped with an 0.1 rad 8-mm diameter cone plate. After co-culture in 12 well plates, BUD-ELMs were directly loaded onto the instrument for analysis. To prevent drying, approximately 100 µL of mineral oil was applied around the circumference of the cone-plate geometry after loading. Strain sweep experiments from 0.1 to 100% strain amplitudes were performed at a fixed frequency of 3.14 rad/s. Frequency sweep experiments from 100 to 0.1 rad/s were performed at a 0.35% strain amplitude. Data were acquired using TRIOS software (version 4.2.1.36612).

## Code Availability

All code generated for this study will be provided via GitHub upon complete submission to a journal, or upon request.

## Acknowledgements

We thank Dr. Elton P. Hudson for graciously providing the wild-type *Synechocystis* strain used in this study, and Dr. Shyam Bhakta for providing DNA parts in the construction of the PilA1 plasmids. This work was primarily supported by the Cancer Prevention and Research Institute of Texas (RR190063, C.M.A.-F.). Additional support was provided by the Welch Foundation (C-2130-20230405), the National Science Foundation Research Traineeship Program, and the GEM Fellowship program.

## Author Contributions

Conceptualization: R.F.T. and C.M.A.-F. Data Curation: R.F.T. Formal Analysis: R.F.T. Funding Acquisition: C.M.A.-F. Investigation: R.F.T., N.E.H., and O.D.S. Methodology: R.F.T., O.D.S., and C.M.A.-F. Project Administration: C.M.A.-F. Resources: R.F.T. Software: R.F.T. Supervision: C.M.A.-F. and R.F.T. Validation: R.F.T. Visualization: R.F.T. Writing (Original Draft): R.F.T and C.M,A.-F. All authors contributed to reviewing and editing this manuscript.

